# Development of a rapid and highly accurate method for ^13^C tracer-based metabolomics and its application on a hydrogenotrophic methanogen

**DOI:** 10.1101/2023.08.31.555715

**Authors:** Yuto Fukuyama, Shigeru Shimamura, Sanae Sakai, Yuta Michimori, Tomomi Sumida, Yoshito Chikaraishi, Haruyuki Atomi, Takuro Nunoura

## Abstract

Microfluidic capillary electrophoresis-mass spectrometry (CE-MS) is a rapid and highly accurate method to determine isotopomer patterns in isotopically labeled compounds. Here, we developed a novel method for tracer-based metabolomics using CE-MS for underivatized proteinogenic amino acids. The method consisting of a ZipChip CE system and a high-resolution Orbitrap Fusion Tribrid mass spectrometer allows us to obtain highly accurate data from 1 μL of 100 nmol/L mol amino acids comparable to a mere 1 × 10^4-5^ prokaryotic cells. To validate the capability of the CE-MS method, we analyzed 16 protein-derived amino acids from a methanogenic archaeon *Methanothermobacter thermautotrophicus* as a model organism, and the mass spectra showed sharp peaks with low mass errors and background noise. Tracer-based metabolome analysis was then performed to identify the central carbon metabolism in *M. thermautotrophicus* using ^13^C-labeled substrates. The mass isotopomer distributions of serine, aspartate, and glutamate revealed the co-occurrence of the Wood-Ljungdahl pathway and an incomplete reductive TCA cycle for carbon fixation. In addition, biosynthesis pathways of 15 amino acids were constructed based on the mass isotopomer distributions of the detected protein-derived amino acid, genomic information, and public database. Among them, the presence of the alternative enzymes of alanine dehydrogenase, ornithine cyclodeaminase, and homoserine kinase was suggested in the biosynthesis pathways of alanine, proline, and threonine, respectively. To our knowledge, the novel ^13^C tracer-based metabolomics using CE-MS is the most efficient method to identify central carbon metabolism and amino acid biosynthesis pathways and is applicable to in any kind of isolated microbe.

## Introduction

Metabolomics is an effective method to detect the metabolic states of microbial cells. Intracellular metabolite levels are regulated in a complex manner responding to metabolite generation and/or consumption brought about by enzymatic, non-enzymatic, and transport reactions [1, 2]. For over the two decades, ^13^C tracer analysis has commonly been utilized as an accurate and physiologically reliable method to examine the changes in intracellular metabolite levels (metabolic fluxes) [3]. ^13^C tracer-based metabolomics provide the means to identify metabolic pathways and their direction because the patterns of the detected ^13^C-labeled metabolites reflect the amount of metabolic flux [1]. Based on ^13^C labeling patterns in key metabolites, active pathways can be traced back, and in some cases the presence of new enzymes can be revealed.

To date, several approaches to identify and quantify intermediate metabolites have been examined using gas chromatography (GC) with mass spectrometry (GC-MS), liquid chromatography (LC) with mass spectrometry (LC-MS), or LC with nuclear magnetic resonance (LC-NMR) [4, 5, 6]. Currently, GC-MS, which is more sensitive than the other methods, is the most popular technique to determine isotopologues of amino acids and subsequently identify novel carbon fixation pathways [7, 8]. However, even with the high sensitivity of the GC-MS technique, large biomass grown with ^13^C-labeled substrates is sometimes required [9]. In addition, background noises of the mass spectra from GS-MS data interfere with the identification of ^13^C-labeled compounds and determination of ^13^C distribution. Thus, to overcome these issues, analytical software to interpret GC-MS data has been developed for the isotopomer analysis of ^13^C-labeled amino acids [10]. For instance, MassWorks software and Isotopo software, a formula determination tool, provide spectra accuracy based on exact calibration from relatively small biomass samples [11, 12, 13, 14]. From another point of view, in GC-MS analysis, derivatization must be employed to improve sensitivity and separation. On the other hand, recently, microfluidic capillary electrophoresis coupled to mass spectrometry (CE-MS) has been developed as a simple and rapid technique for metabolomic studies. The CE-MS method does not require abundant biomass nor derivatization of molecules for analysis, and provides data with low background noise [15, 16]. This method can thus be expected to be applicable for tracer-based metabolomics to obtain high-quality data from smaller biomass compared to GC-MS based metabolomics.

*Methanothermobacter thermautotrophicus* is a model organism of thermophilic and hydrogenotrophic methanogens. The methanogen grows chemolithoautotrophically by generating methane using H_2_ and CO_2_ as sole energy and carbon sources, respectively [17, 18, 19]. In this methanogen, CO_2_ is sequentially reduced to methane via the archaeal type Wood-Ljungdahl (WL) pathway for energy conservation and to acetyl-CoA for carbon fixation. In addition, in combination with classic tracer-based metabolomics and genomic information, the capability of carbon fixation with an incomplete reductive TCA cycle in this methanogen has also been proposed [20, 21, 22, 23]. From genomic information, the organism harbors a gene set of an almost complete TCA cycle but lacks aconitase hydratase (EC 4.2.1.3) and isocitrate dehydrogenase (EC 1.1.1.42) [22]. The metabolic function of the putative incomplete TCA cycle has been partly revealed by several radioisotope tracer analyses [20, 21]. The methanogen also incorporated acetate as a carbon source via reductive operation of the TCA cycle via acetyl-CoA carboxylation under physiological conditions [21, 24] despite the absence of acetogenotrophic metabolism [17]. However, experimental evidence is limited to verify the metabolic function of the central carbon metabolism of *M. thermautotrophicus* and the roles of the incomplete TCA cycle.

Here, we developed a novel, rapid, sensitive, and highly accurate tracer-based metabolomics method for proteinogenic amino acids using CE-MS and CE-MS/MS. The analytical method requires only 1 μL of 100 nmol/L amino acids that correspond to those from approximately 10^4-5^ prokaryotic cells. Total run time, including wash, was 15 minutes for each sample. Moreover, the method reveals the position of labeled carbon using CE-MS/MS analysis, which is essential to understand the carbon metabolic pathways and amino acid biosynthesis pathways. In this study, the capability of the incomplete reverse TCA cycle (rTCA) in *M. thermautotrophicus* was examined based on the isotopologues and isotopomer analysis as the first ^13^C tracer-based metabolomics report using the ZipChip CE system and the high-resolution Orbitrap Fusion Tribrid mass spectrometer. Furthermore, all amino acid biosynthesis pathways were reconstructed. A dataset of the ^13^C tracer-based metabolomics together with genomic information provides the most probable central carbon metabolism in this methanogen along with its amino acid biosynthesis pathways. The novel CE-MS and CE-MS/MS techniques is a promising tool to reveal the amino acid biosynthesis pathways and related central carbon metabolism that are essential to understand the core anabolism of life in phylogenetically and metabolically diverse organisms including those that are not applicable for metabolomics because of the difficulties to obtain sufficient biomass.

## Materials and Methods

### Strain and growth conditions

*Methanothermobacter thermautotrophicus* delta H^T^ (=JCM10044^T^) was obtained from the Japan Collection of Microorganisms (JCM) in Riken. Unless otherwise indicated, *M. thermautotrophicus* was grown with a 3 mL JCM231 medium in 17 mL test tubes at 65°C. Headspace gas was balanced to atmospheric pressure with H_2_/CO_2_ (80:20, v/v).

### *In vivo incorporation* of ^13^C-labeled substrates

All the ^13^C-labeled reagents used in this study were purchased from Cambridge isotope laboratories (CIL) (Tewksbury, MA, USA). For tracer-based metabolomics, *M. thermautotrophicus* was grown in JCM231 medium with ^13^C-labeled substrates as follows. After 24 hours of inoculation, 0.14 mL ^13^CO_2_ was added to the headspace of the culture. [1-^13^C_1_] or [2-^13^C_1_] sodium acetate was added in JCM231 medium (final concentration 0.01% (v/v)) before inoculation. Cells in the three mL of the culture were harvested at the exponential phase by centrifugation (4°C, 10,000 × g, 10 min), frozen in liquid N_2_, and then stored at -80°C until use. Negative control cultures were also prepared under the same conditions in the absence of ^13^C-labeled substrates. All isotope tracing experiments were performed in duplicate.

### Sample preparation

The stored cells were hydrolyzed to prepare protein-derived amino acids in 12 N HCl at 110°C for 16 h in a 1.0 mL glass reaction vial (GL Science, Tokyo, Japan). After adding *n*-hexane to remove hydrophobic constituents, the hydrolysate was filtered through Nanosep MF centrifugal Devices with 0.45 μm wwPTFE membrane (PALL, Port Washington, NY, USA) by centrifugation. The filtered sample was washed with *n*-hexane/dichloromethane (1:2, v/v) three times. The dried protein-derived amino acids were obtained with evaporation under an N_2_ stream and then stored at -80°C until use.

### Isotopologue and isotopomer analyses

Isotope abundance and incorporation rate were examined using a ZipChip CE system coupled with an Orbitrap Fusion Tribrid mass spectrometer (Thermo Fisher Scientific, Waltham, MA, USA). The CE separation was performed with an HS chip (908 Devices, Boston, MA, USA) in the following manner: injection volume 5 nL, field strength 1,000 V/cm, analysis run time 3 minutes without pressure assist. The MS analysis was operated in positive ion mode to detect targeted proteinogenic amino acids based on their accurate masses. The full scan MS settings included: Detector Orbitrap, Resolution 15,000, Quadrupole Isolation ON, Scan Range 70-500 *m/z*, an acquisition gain control (AGC) Target Standard, Maximum Injection Time Mode 50 ms, EASY-IC ON, Sheath Gas Flow Rate 2.2 arbitrary units (au), Capillary Temperature at 200°C, S-lens Radio Frequency (RF) Level 60%. The MS/MS settings included: Isolation Mode quadrupole, Isolation Window 0.7 *m/z*, Activation Type HCD, Collision Energy Mode stepped (25, 35, 50), Detector Type Orbitrap, Resolution 15,000, Scan Range Mode Define First Mass (*m/z* 40), AGC Target Standard, Maximum Injection Time Mode Dynamic. The obtained data was analyzed using Qual Browser in Xcalibur version 4.3.73.11 (Thermo Fisher Scientific). Prior to the analysis for the protein-derived amino acids, a complete standard mixture of amino acids (Promega, Medison, WI, USA) was analyzed by CE-MS to obtain MS and MS/MS spectra of the proteinogenic amino acids. Non-labeled, [1-^13^C_1_], [5-^13^C_1_], and [^13^C_5_] glutamate were also examined for validation of the capability of MS/MS fragmentation. Detected protein-derived amino acids were identified by migration times and monoisotopic mass matching to the amino acid mixture under the same condition. ^13^C-labeled positions and their ^13^C abundance were estimated by fragment patterns based on ^13^C-labeled standard of amino acids and an MS spectra library (mzCloud) (Thermo Fisher Scientific).

To identify the central carbon metabolism and amino acid biosynthesis pathways based on the isotopologue patterns of amino acids, we compared the observed mass isotopomer distributions with predicted labeling structures from genomic information in KEGG PATHWAY Database [25] and MetaCyc [26]. Manual curation of amino acid biosynthesis pathways was performed by a web-tool, GapMind [27].

## Results and discussion

### Identification of amino-acids using the CE-MS method

To examine the capability of the CE-MS method for identification of amino acids, we analyzed a standard mixture of amino acids. All amino acids are positively charged in this method using an acidic running buffer and are thus separated in response to the differences in electrophoretic mobility based on each side chain structure. Accordingly, positively charged amino acids are identified in shorter migration times followed by neutral amino acids and then the acidic amino acids (Fig. 1A). We next analyzed 20 protein-derived amino acids from *M. thermautotrophicus* cells grown with non-labeled substrate (Fig. 1B). Sharp peaks of 16 proteinogenic amino acids were identified with slight background noise in a short run (3 minutes) while those of cysteine, tryptophan, asparagine, and glutamine were not detected. It has been pointed out that cysteine and tryptophan are degraded during hydrolyzation by 12 N HCl at high temperature [9, 28]. In addition, asparagine and glutamine are converted to aspartate and glutamate, respectively, in 12 N HCl [9, 28]. The mass spectra of the detected cellular protein-derived amino acids were consistent with those from the standard amino acid mixture. Mass errors ranged from 0.04 to 4.61 ppm when we compared the theoretical *m/z* and an experimentally observed *m/z*. Peak areas of the detected amino acids ranged from 4,119 to 20,448,890. The lower limit of this method was 1 μL of 100 nmol/L mol amino acids that correspond with amino acids from approximately 10^4-5^ prokaryotic cells. These results indicated high accuracy and sensitivity of the CE-MS method. <u>Isotopologue characterization using the CE-MS/MS method</u> To validate the applicability of the developed CE-MS/MS method for isotopomer analysis, we analyzed MS and MS/MS spectra of non-labeled, [1-^13^C_1_], [5-^13^C_1_], and [^13^C_5_] glutamate (Fig 2). The observed labeled fragments in positive ionization were consistent with the numbers and localization of the labeled carbons in each glutamate. In conventional tracer-based metabolomics using GC-MS, high accuracy data are adjusted by an analytical software to distinguish ^13^C distribution in each derivatized amino acid. By contrast, the developed CE/MS and CE-MS/MS methods provided sufficiently clear and accurate MS/MS spectra without complicated sample pretreatment. Thus, we concluded that the developed method was applicable in providing practical clues to characterize isotopologues of amino acids.

**Fig. 1.**
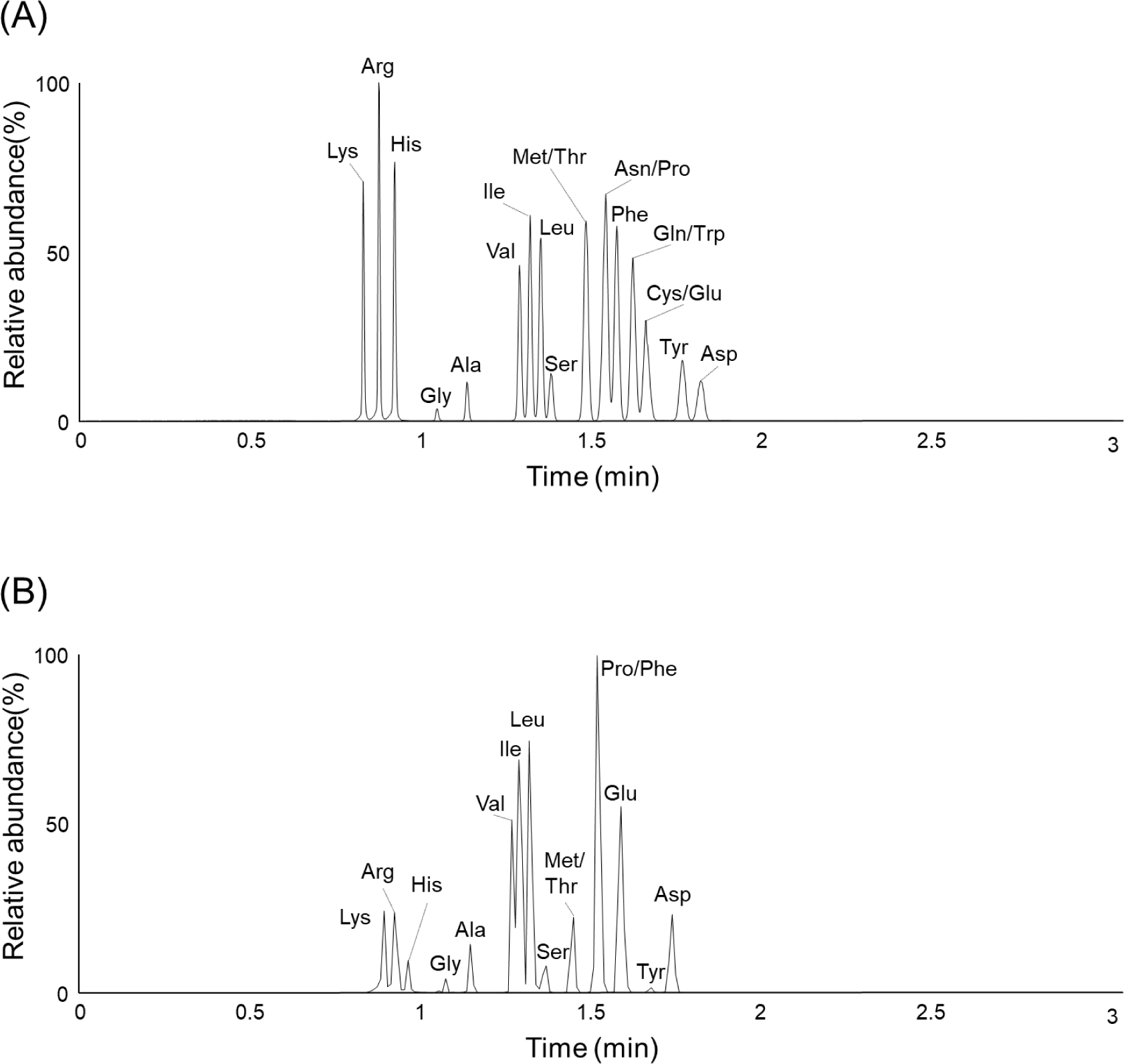
Mass spectra of the detected amino acids from the standard mixture of amino acids (A) and *M. thermautotrophicus* (B) Gly glycine, Ala alanine, Ser serine, Thr threonine, Asn asparagine, Gln glutamine, Asp aspartate, Glu glutamate, Lys lysine, Arg arginine, His histidine, Val valine, Leu leucine, Ile isoleucine, Tyr tyrosine, Phe phenylalanine, Trp tryptophan, Pro proline, Met methionine, Cys cysteine.

**Fig. 2.**
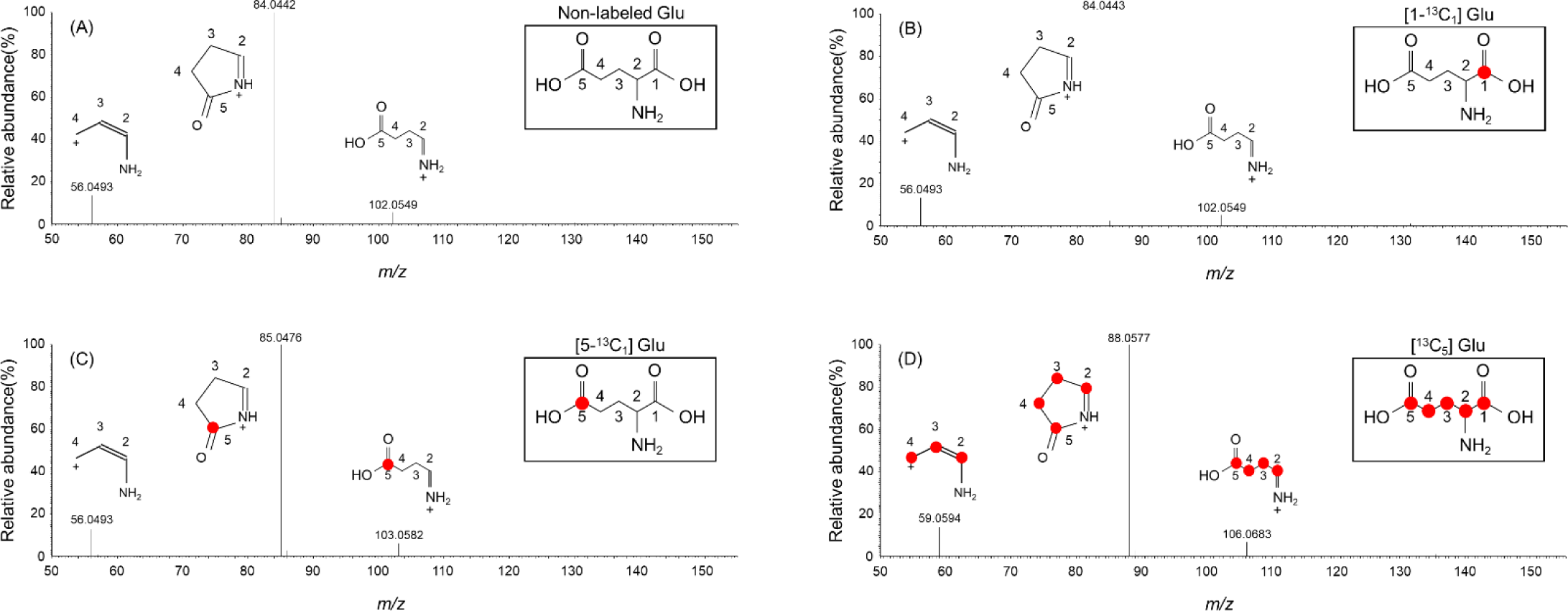
MS/MS spectra of non-labeled glutamate (A), [1-^13^C_1_] glutamate (B), [5-^13^C_1_] glutamate (C), and [^13^C_5_] glutamate (D) Red circles show ^13^C in the carbon skeleton.

### Co-occurrence of the WL pathway and the incomplete reductive TCA cycle in *M. thermautotrophicus*

To verify the central carbon metabolism and the amino acid biosynthesis pathways in *M. thermautotrophicus*, we applied the CE-MS/MS method for isotopomer analysis. The methanogen is expected to harbor the WL pathway and an incomplete rTCA cycle for carbon fixation based on genomic analyses [22, 23]. Genomic information suggests that serine (Ser), aspartate (Asp), and glutamate (Glu) are synthesized from pyruvate, oxaloacetate, and 2-oxoglutarate, respectively (Fig. 3). In this study, ^13^CO_2_ and ^13^C labeled acetate (final concentration, 0.01 % wt/v) were used as a tracer to identify the carbon fixation route of the methanogen. The low concentration of the ^13^C labeled acetate was expected not to affect the metabolic flux of the inorganic carbon fixation pathways.

**Fig. 3.**
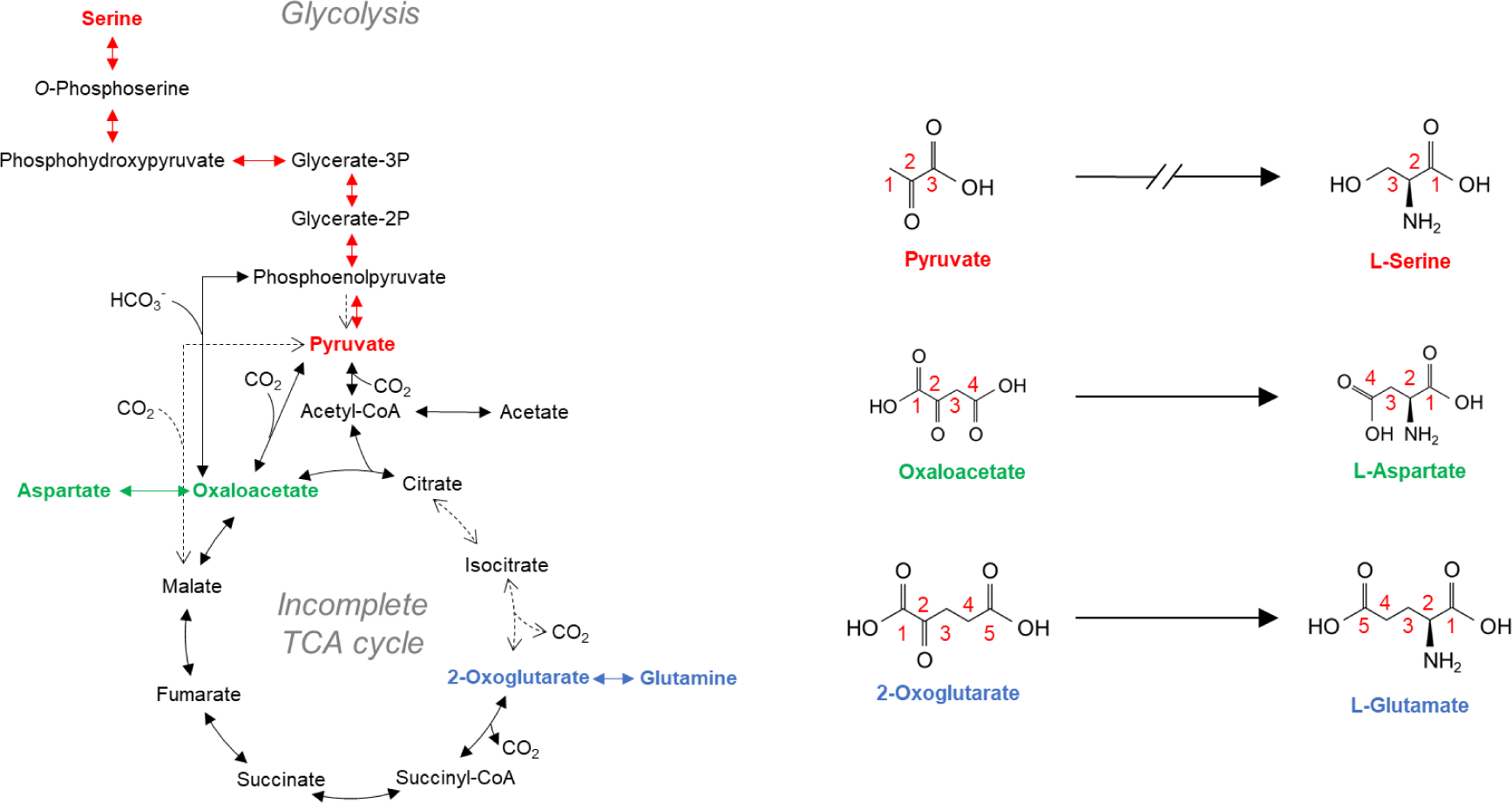
Carbon transitions of serine, aspartate, and glutamate via each biosynthesis pathway based on genomic information. Numbers in each structural formula show carbon numbers.

In cells grown with non-labeled substrates, isotopomers labeled with natural ^13^C were detected in the mass fractions of M+1 (Fig. 4, Table 1). The relative abundance of the mass fraction labeled with natural ^13^CO_2_ (M+1 fraction) in Ser, Asp, and Glu varied from each other. The result suggested that the incorporation of natural ^13^CO_2_ affected the isotopomers composition of the precursors of Ser, Asp, and Glu. Thus, the incorporation rate of ^13^C *in vivo* labeling was evaluated after eliminating the natural abundance of ^13^C (1.1%) (Table 1). In cells grown with ^13^CO_2_, up to three labeled carbons were detected in Ser (Fig. 4 and Table 1), and ^13^C incorporation was found in both main-and side-chains (Table 1). The isotopomer patterns were consistent with the known carbon fixation via the WL pathway [22], followed by carboxylation of acetyl-CoA to form pyruvate and by serine biosynthesis from pyruvate via 3-phosphoglycerate. Similarly, Asp harbored up to three labeled carbons (Fig. 4 and Table 1). The ^13^C incorporation in the side-chain of Asp is explained by an anaplerotic reaction (malate shunt), which synthesizes oxaloacetate (the precursor of Asp) from pyruvate and CO_2_, and subsequent Asp biosynthesis from oxaloacetate (Fig. 5A). Glu also had up to three labeled carbons (Fig. 4 and Table 1). The ^13^C incorporation at the main-chain of Glu can be explained by carboxylation of succinyl-CoA to form 2-oxoglutarate (Fig. 5A), and subsequent Glu biosynthesis from 2-oxoglutarate. If a complete reductive TCA cycle (rTCA) occurred, completely labeled Asp and Glu should be detected in cells grown with ^13^CO_2_ due to multiple rounds of conversion through the rTCA cycle as reported previously [14]

**Fig. 4.**
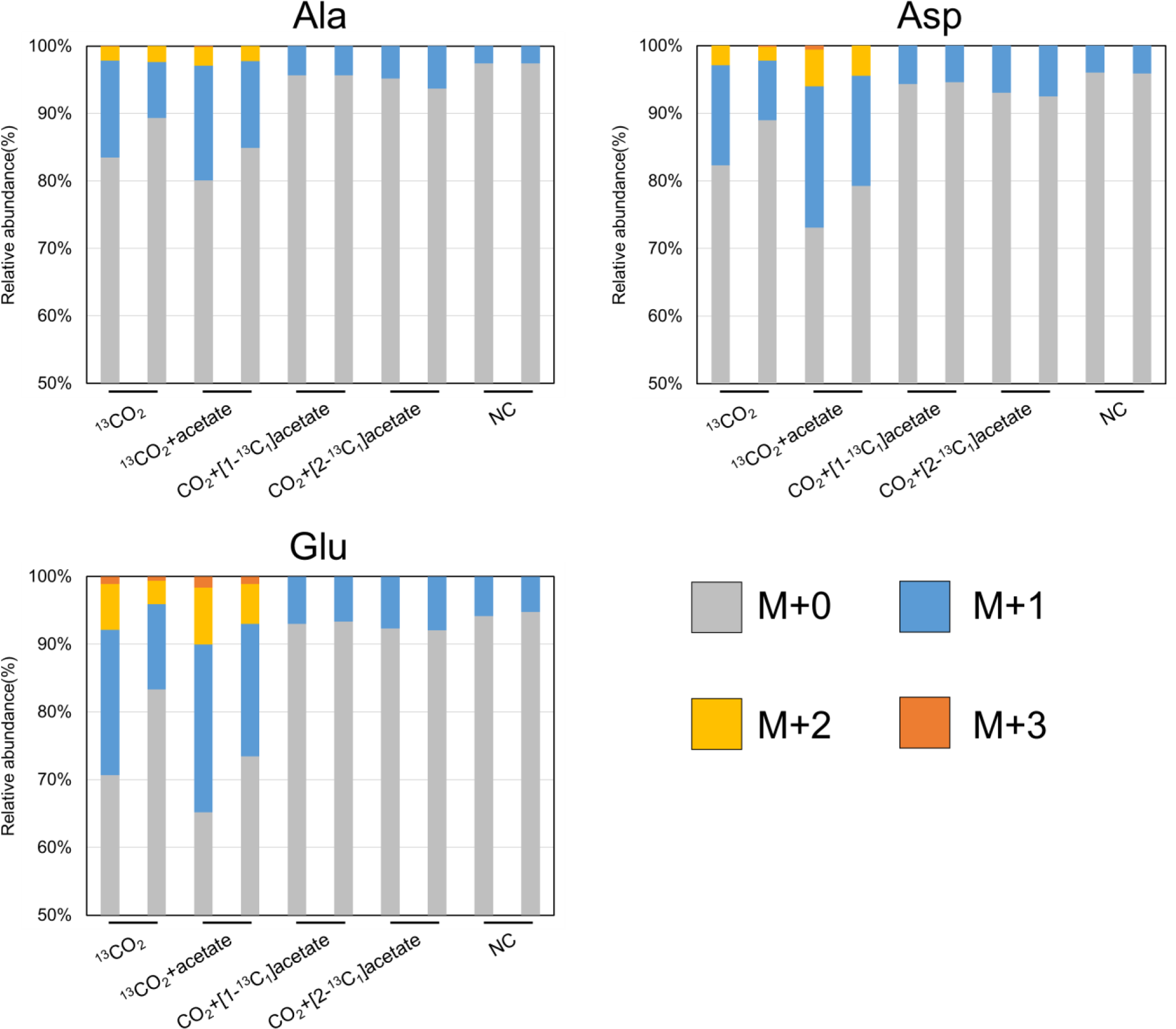
Mass isotopomer distributions from selected amino acids for the WL pathway, the incomplete TCA cycle, and gluconeogenesis. The mass fractions for M+0, M+1, M+2, and M+3, which represent fragments containing 0 to 3 ^13^C-labeled carbons, respectively.

**Table 1.**
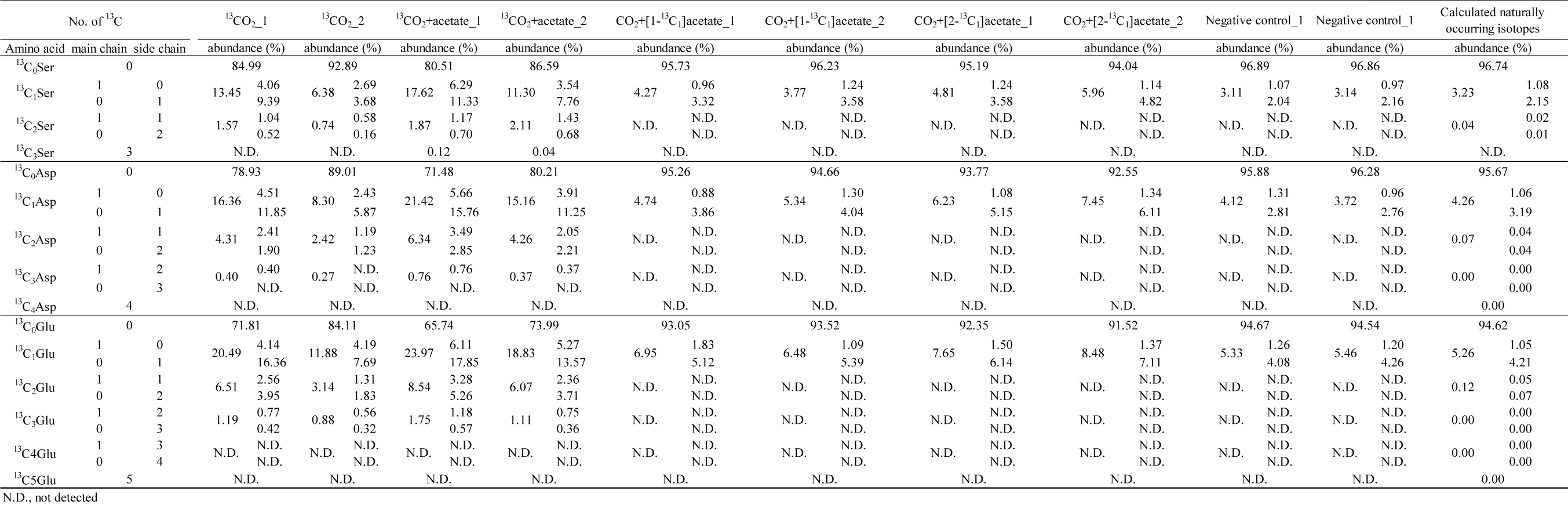
The number and abundance of ^13^C in serine, aspartate, and glutamate from the isotopomer analysis.

**Fig. 5.**
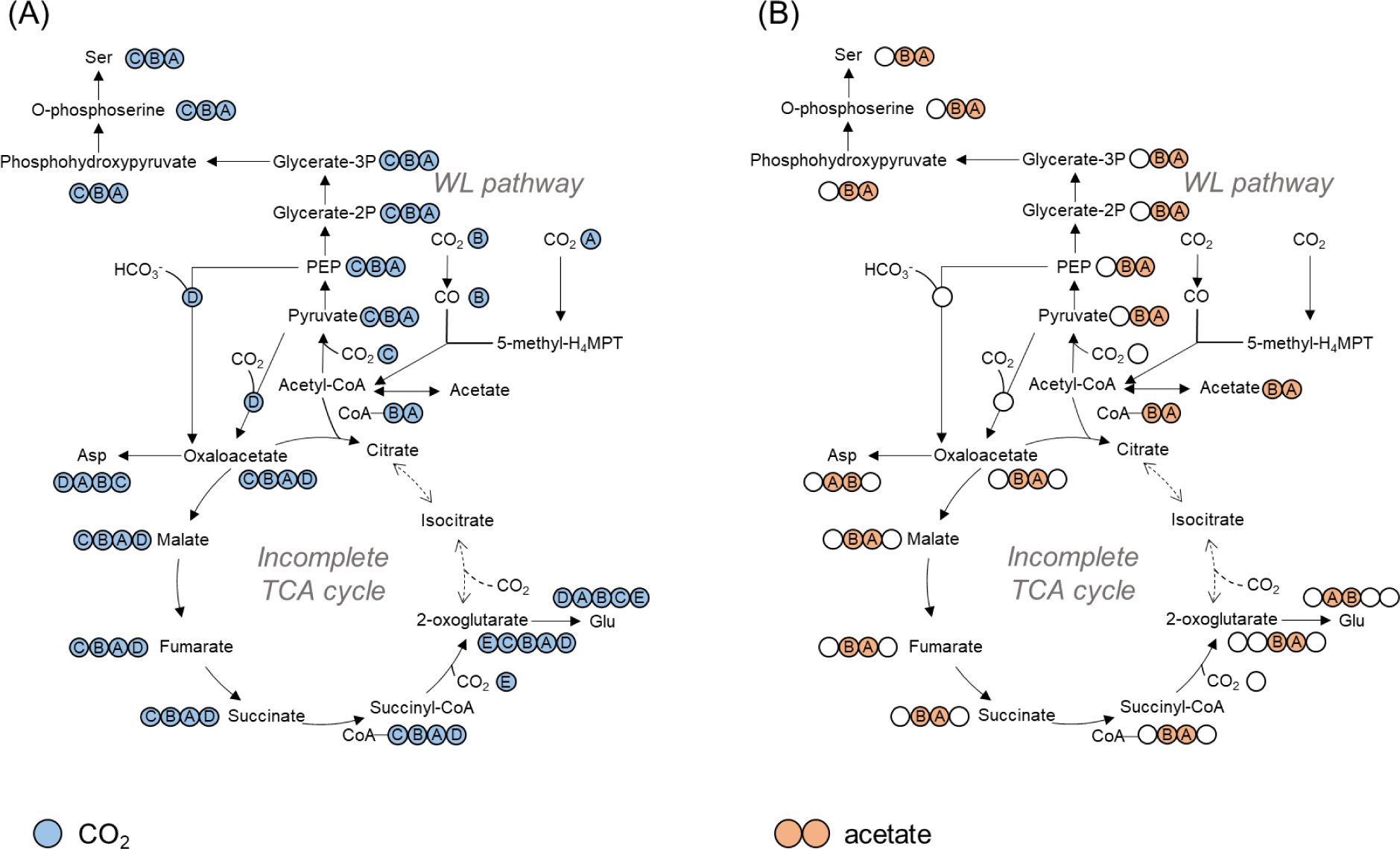
Elucidating the co-occurrence of the WL pathway and the incomplete reductive TCA cycle. Proposed labelling of metabolites from metabolome analysis with ^13^CO_2_ (A) and ^13^C-labeled acetate ([1-^13^C_1_] or [2-^13^C_1_] acetate) (B) during the WL pathway and incomplete TCA cycle. Circles indicate carbon atoms in the metabolites. Dot lines indicate there are no corresponding enzymes found in the genome.

We also examined the isotopomer analysis using ^13^C labeled acetate; [1-^13^C_1_] or [2-^13^C_1_] acetate (0.01 % wt/v). The ^13^C incorporation pattern showed that the relative abundance of mass fraction of M+1 in each amino acid increased compared to those in the negative control (Fig. 4 and Table 1). In addition, the labeled carbon was also observed in the side-chain of each amino acid (Table 1). These results demonstrated that acetate was converted to acetyl-CoA, and incorporated into the TCA cycle under reductive operation but the cycle was incomplete as suggested by the ^13^CO_2_ tracer experiment (Fig. 5). Accordingly, we conclude that our data set strongly supports the co-occurrence of the WL pathway and an incomplete rTCA cycle in *M. thermautotrophicus*, that was suggested by the absence of genes for aconitase and isocitrate dehydrogenase [22, 23] and the previously reported ^14^C tracer-based metabolomics [20, 21]. The conclusion is consistent with a recent kinetic carbon network analysis whose results suggest that acetyl-CoA influx from the WL pathway and/or another central carbon metabolic pathway yielding acetyl-CoA conflicts with the complete rTCA cycle [29].

### Amino acid biosynthesis pathways of *M. thermautotrophicus*

The KEGG PATHWAY database is a visual collection of metabolic pathways and one of the most commonly used bioinformatic resources to identify enzymatic reactions and metabolic functions from genomic perspectives [25, 30]. The complete genome of *M. thermautotrophicus* in the KEGG PATHWAY database suggests that a number of reactions/enzymes in several amino acid biosynthesis pathways, including that for proline, cannot be clearly defined. The amino acid biosynthesis pathways obtained from the KEGG PATHWAY database were compared against the MetaCyc database, the most extensively curated collection of metabolic pathways containing 2749 pathways derived from more than 60,000 publications [26]. Then, GapMind [27] was used to curate the amino acid biosynthesis pathways and to identify candidate enzymes that were not present in the KEGG pathway (*M. thermautotrophicus*). GapMind is a web-based tool for annotating amino acid biosynthesis pathways in bacteria and archaea, where each curated pathway is supported by a confidence level [27]. However, three metabolic gaps in the amino acid biosynthesis pathways of *M. thermautotrophicus* still remained; genes for alanine transaminase (EC 2.6.1.2), ornithine cyclodeaminase (EC 4.3.1.12), and homoserine kinase (EC 2.7.1.39), in the most probable alanine, proline and threonine biosynthesis pathways, respectively.

To examine the probability of the amino acid biosynthesis pathways of *M. thermautotrophicus* in the KEGG PATHWAY database and MetaCyc, the mass isotopomer distributions of the detected 16 protein-derived amino acids in this methanogen were compared with the expected isotopomer structures figured out by GapMind (Fig. S1 and Table S1). In this analysis, glycine was excluded because the *m/z* value of fragmented glycine was below the detection limit. In addition, a histidine biosynthesis pathway could not be confirmed because the relative abundance of MS/MS spectra from fragmented histidine was not sufficient in cells labeled with [1-^13^C_1_] acetate or [2-^13^C_1_] acetate (Table S1). All the amino acids detected in this study harbored one or more ^13^C (Table S1). The labeling pattern of each amino acid was interpreted by the predicted amino acids biosynthesis pathways constructed from a combination of the public databases (KEGG PATHWAY and MetaCyc databases) and previously reported tracer-based metabolomics using radioisotope including proline, threonine, and alanine biosynthesis pathways [31, 32, 33] (Fig. S1). For example, the isotopomer patterns of Ser and aromatic amino acids suggested that pyruvate was their common precursor and supplied via gluconeogenesis (Fig. S1 and Table S1).

Based on the isotopomer analysis, we also explored alternative enzymes to fill the metabolic gaps in alanine, proline, threonine, and alanine biosynthesis pathways again. The isotopomer pattern of alanine was similar to that of serine, suggesting that alanine was synthesized from pyruvate by an alternative enzyme for a determination from pyruvate to alanine. Generally, this deamination is reversibly catalyzed by alanine dehydrogenase (EC 1.4.1.1) or alanine transaminase (EC 2.6.1.2). In the *M. thermautotrophicus* genome, a candidate gene (MTH_1495) showed significant amino acid sequence identity with a newly characterized archaeal alanine dehydrogenase in *Archaeoglobus fulgidus* [34, 35]. On the other hand, when we searched for subgroup I transaminase, in which alanine transaminase was included [36], three genes (MTH_1894, MTH_1694, and MTH_52) were identified in their genome. Of these genes, MTH_1894 and MTH_1694 were annotated as aspartate aminotransferase (EC 2.6.1.1), and MTH_52 was annotated as LL-diaminopimelate aminotransferase (EC 2.6.1.83). Transaminases often recognize a broad range of substrates and predicting their specific substrate is challenging [37]. Thus, the archaeal alanine dehydrogenase and the three subgroup I transaminases in *M. thermautotrophicus* genome could be candidates for alternative enzyme for the reversible deamination between pyruvate and alanine. Similarly, in the case of proline and threonine, the observed isotopomer structures were consistent with the most probable biosynthesis pathways including previously unknown ornithine cyclodeaminase and homoserine kinase, respectively (Fig. S1). Recently, it was reported that an archaeal ornithine cyclodeaminase (MMP_1218) in *Methanococcus maripaludis* produced proline from ornithine [38]. In *M. thermautotrophicus*, MTH_867 showed significant amino acid sequence identity with MMP_1218 (58.75%), suggesting that archaeal ornithine cyclodeaminase encoded by MTH_867 filled the metabolic gap in the proline biosynthesis pathway. However, no possible alternative enzymes were reported for the metabolic gap in the threonine biosynthesis pathway, and similar metabolic gaps are also observed in other microbes, such as *Methanocaldococcus jannaschii*, *Bacteroides thetaiotaomicron, Dinoroseobacter shibae*, and *Phaeobacter inhibens* [37].

The prediction of the amino acid biosynthesis pathways in public databases is very useful to understand the metabolic and genetic capability of organism including secondary metabolites. However, the limitations of the databases have been pointed out because these are based on biochemical, molecular biological, and genomic information from a limited number of model organisms, and thus, alternative biosynthesis pathways are often missing [37, 39]. The developed ^13^C tracer-based metabolomics using ZipChip CE system in combination with Orbitrap Fusion Tribrid mass spectrometer can help in predicting the most probable metabolic pathway and provide clues to reveal the presence of unknown alternative enzymes or unknown amino acid biosynthesis pathways that are not recognized in the public database. The developed tracer-based metabolomic method for the amino acids is a promising tool to identify metabolic pathways related to amino acids from any kind of isolated microbes due to its high sensitivity and accuracy conveniently, and contribute to construct more reliable databases of microbial metabolic pathways.

## Acknowledgments

A part of this work was supported by Grant-in-Aid for Scientific Research on Innovative Areas “Post-Koch Ecology” (19H05684) and (A) (19H00988) from The Ministry of Education, Culture, Sports, Science and Technology (MEXT) and the Green Innovation Fund Project, JPNP22010, commissioned by the New Energy and Industrial Technology Development Organization (NEDO).

## References

1. Buescher JM, Antoniewicz MR, Boros LG, Burgess SC, Brunengraber H, Clish CB, et al. A roadmap for interpreting ^13^C metabolite labeling patterns from cells. Curr Opin Biotechnol. 2015; 34: 189–201. 10.1016/j.copbio.2015.02.003

2. Keller MA, Piedrafita G, Ralser M. The widespread role of non-enzymatic reactions in cellular metabolism. Curr Opin Biotechnol. 2015; 34: 153–161. 10.1016/j.copbio.2014.12.020

3. Crown SB, Antoniewicz MR. Publishing ^13^C metabolic flux analysis studies: A review and future perspectives. Metab Eng. 2013; 20: 42–48. 10.1016/j.ymben.2013.08.005

4. Mohabbat T, Drew B. Simultaneous determination of 33 amino acids and dipeptides in spent cell culture media by gas chromatography-flame ionization detection following liquid and solid phase extraction. J Chromatogr B. 2008; 862: 86–92. 10.1016/j.jchromb.2007.11.003

5. Le A, Ng A, Kwan T, Cusmano-Ozog K, Cowan TM. A rapid, sensitive method for quantitative analysis of underivatized amino acids by liquid chromatography–tandem mass spectrometry (LC-MS/MS). J Chromatogr B. 2014; 944: 166–174. 10.1016/j.jchromb.2013.11.017

6. Villas-Bôas SG, Mas S, Åkesson M, Smedsgaard J, Nielsen J. Mass spectrometry in metabolome analysis. Mass Spectrom Rev. 2005; 24: 613–646. 10.1002/mas.20032

7. Huber H, Gallenberger M, Jahn U, Eylert E, Berg IA, Kockelkorn D, et al. A dicarboxylate/4-hydroxybutyrate autotrophic carbon assimilation cycle in the hyperthermophilic Archaeum *Ignicoccus hospitalis*. Proc Natl Acad Sci U S A. 2008; 105: 7851–7856. 10.1073/pnas.0801043105

8. Xiong W, Lo J, Chou KJ, Wu C, Magnusson L, Dong T, et al. Isotope-assisted metabolite analysis sheds light on central carbon metabolism of a model Cellulolytic bacterium *Clostridium thermocellum*. Front Microbiol. 2018; 9: 1947. 10.3389/fmicb.2018.01947

9. Tang YJ, Martin HG, Myers S, Rodriguez S, Baidoo EEK, Keasling JD. Advances in analysis of microbial metabolic fluxes via ^13^C isotopic labeling. Mass Spectrom Rev. 2009; 28: 362–375. 10.1002/mas.20191

10. Peterson AC, McAlister GC, Quarmby ST, Griep-Raming J, Coon JJ. Development and characterization of a GC-enabled QLT-orbitrap for high-resolution and high-mass accuracy GC/MS. Anal Chem. 2010; 82: 8618–8628. 10.1021/ac101757m

11. Misra, BB. Advances in high resolution GC-MS technology: a focus on the application of GC-Orbitrap-MS in metabolomics and exposomics for FAIR practices. Analytical Methods. 2021; 13: 2265–2282. 10.1039/D1AY00173F

12. Nunoura T, Chikaraishi Y, Izaki R, Suwa T, Sato T, Harada T, et al. A primordial and reversible TCA cycle in a facultatively chemolithoautotrophic thermophile. Science. 2018; 359:559–563. 10.1126/science.aao3407

13. Ahmed Z, Zeeshan S, Huber C, Hensel M, Schomburg D, Münch R, et al. ‘Isotopo’ a database application for facile analysis and management of mass isotopomer data. Database. 2014; 2014: bau3 10.1093/database/bau077

14. Steffens L, Pettinato E, Steiner TM, Mall A, König S, Eisenreich W, et al. High CO_2_ levels drive the TCA cycle backwards towards autotrophy. Nature. 2021; 592: 784–788. 10.1038/s41586-021-03456-9

15. Zhang W, Ramautar R. CE-MS for metabolomics: Developments and applications in the period 2018–2020. Electrophoresis. 2021; 42: 381–401. 10.1002/elps.202000203

16. Ribeiro da Silva M, Zaborowska I, Carillo S, Bones J. A rapid, simple and sensitive microfluidic chip electrophoresis mass spectrometry method for monitoring amino acids in cell culture media. J of Chromatogr A. 2021; 1651: 462336. 10.1016/j.chroma.2021.462336

17. Zeikus JG, Wolfe RS. *Methanobacterium thermoautotrophicus* sp. n., an Anaerobic, Autotrophic, Extreme Thermophile. J Bacteriol. 1972; 109: 707–715. 10.1128/jb.109.2.707-713.1972

18. Thauer RK, Kaster AK, Seedorf H, Buckel W, Hedderich R. Methanogenic archaea: ecologically relevant differences in energy conservation. Nat Rev Microbiol. 2008; 6: 579–591. 10.1038/nrmicro1931

19. Conrad R. The global methane cycle: recent advances in understanding the microbial processes involved. Environ Microbiol Rep. 2009; 1: 285–292.

20. Fuchs G, Stupperich E. Evidence for an incomplete reductive carboxylic acid cycle in *Methanobacterium thermoautotrophicum*. Arch Microbiol. 1978; 118: 121–125. 10.1111/j.1758-2229.2009.00038.x

21. Fuchs G, Stupperich E, Thauer RK. Acetate assimilation and the synthesis of alanine, aspartate and glutamate in *Methanobacterium thermoautotrophicum*. Arch Microbiol. 1978; 117: 61–66. 10.1007/BF00689352

22. Smith DR, Doucette-Stamm LA, Deloughery C, Lee H, Dubois J, Aldredge T, et al. Complete genome sequence of *Methanobacterium thermoautotrophicum* deltaH: functional analysis and comparative genomics. J Bacteriol. 1997; 179: 7135–7155. 10.1128/jb.179.22.7135-7155.1997

23. Makarova KS, Koonin E V. Filling a gap in the central metabolism of archaea: prediction of a novel aconitase by comparative-genomic analysis. FEMS Microbiol Lett. 2003; 227: 17–23. 10.1016/S0378-1097(03)00596-2

24. Oberlies G, Fuchs G, Thauer RK. Acetate thiokinase and the assimilation of acetate in *Methanobacterium thermoautotrophicum*. Arch Microbiol. 1980; 128: 248–252. 10.1007/BF00406167

25. Kanehisa M, Sato Y, Kawashima M, Furumichi M, Tanabe M. KEGG as a reference resource for gene and protein annotation. Nucleic Acids Res. 2016; 44: 457–462. 10.1093/nar/gkv1070

26. Caspi R, Billington R, Keseler IM, Kothari A, Krummenacker M, Midford PE, et al. The MetaCyc database of metabolic pathways and enzymes - a 2019 update. Nucleic Acids Res. 2020; 48: 445–453. 10.1093/nar/gkz862

27. Price MN, Deutschbauer AM, Arkin AP. GapMind: Automated Annotation of Amino Acid Biosynthesis. mSystems. 2020; 5: e00291–20. 10.1128/msystems.00291-20

28. Dauner M, Sauer U. GC-MS Analysis of Amino Acids Rapidly Provides Rich Information for Isotopomer Balancing. Biotechnol Prog. 2000; 16: 642–649. 10.1021/bp000058h

29. Sumi T, Harada K. Kinetics of the ancestral carbon metabolism pathways in deep-branching bacteria and archaea. Comm Chem. 2021; 4: 1–9. 10.1038/s42004-021-00585-0

30. Tanabe M, Kanehisa M. Using the KEGG database resource. Curr Protoc Bioinformatics. 2012; 38: 1.12.1–1.12.43 10.1002/0471250953.bi0112s38

31. Eikmanns B, Linder D, Thauer RK. Unusual pathway of isoleucine biosynthesis in *Methanobacterium thermoautotrophicum*. Arch Microbiol. 1983; 136: 111–113. 10.1007/BF00404783

32. Eisenreich W, Schwarzkopf B, Bacher A. Biosynthesis of nucleotides, flavins, and deazaflavins in *Methanobacterium thermoautotrophicum*. J Biol Chem. 1991; 266: 9622–9631. 10.1016/S0021-9258(18)92866-8

33. Schröder I, Thauer RK. Methylcobalamin:homocysteine methyltransferase from *Methanobacterium thermoautotrophicum*. Eur J Biochem. 1999; 263: 789–796. 10.1046/j.1432-1327.1999.00559.x

34. Schröder I, Vadas A, Johnson E, Lim S, Monbouquette HG. A novel archaeal alanine dehydrogenase homologous to ornithine cyclodeaminase and μ-crystallin. J Bacteriol. 2004; 186: 7680–7689. 10.1128/jb.186.22.7680-7689.20

35. Gallagher DT, Monbouquette HG, Schröder I, Robinson H, Holden MJ, Smith NN. Structure of Alanine Dehydrogenase from *Archaeoglobus*: Active Site Analysis and Relation to Bacterial Cyclodeaminases and Mammalian mu Crystallin. J Mol Biol. 2004; 342: 119–130. 10.1016/j.jmb.2004.06.090

36. Jensen, Roy A., and Wei Gu. Evolutionary recruitment of biochemically specialized subdivisions of family I within the protein superfamily of aminotransferases. J Bacteriol. 1996; 178: 2161–2171. 10.1128/jb.178.8.2161-2171.1996

37. Price MN, Zane GM, Kuehl J V., Melnyk RA, Wall JD, Deutschbauer AM, et al. Filling gaps in bacterial amino acid biosynthesis pathways with high-throughput genetics. PLoS Genet. 2018; 14: e1007147. 10.1371/journal.pgen.1007147

38. Burnat M, Picossi S, Valladares A, Herrero A, Flores E. Catabolic pathway of arginine in *Anabaena* involves a novel bifunctional enzyme that produces proline from arginine. Mol Microbiol. 2019; 111: 883–897. 10.1111/mmi.14203

39. Shrestha P, Kim MS, Elbasani E, Kim JD, Oh TJ. Prediction of trehalose-metabolic pathway and comparative analysis of KEGG, MetaCyc, and RAST databases based on complete genome of *Variovorax* sp. PAMC28711. BMC Genom Data 2022; 23: 1–7. 10.1186/s12863-021-01020-y

