## Supplementary Figure 1 for "Development of a rapid and highly accurate method for ^13^C tracer-based metabolomics and its application on a hydrogenotrophic methanogen"

### Glycolysis / Gluconeogenesis

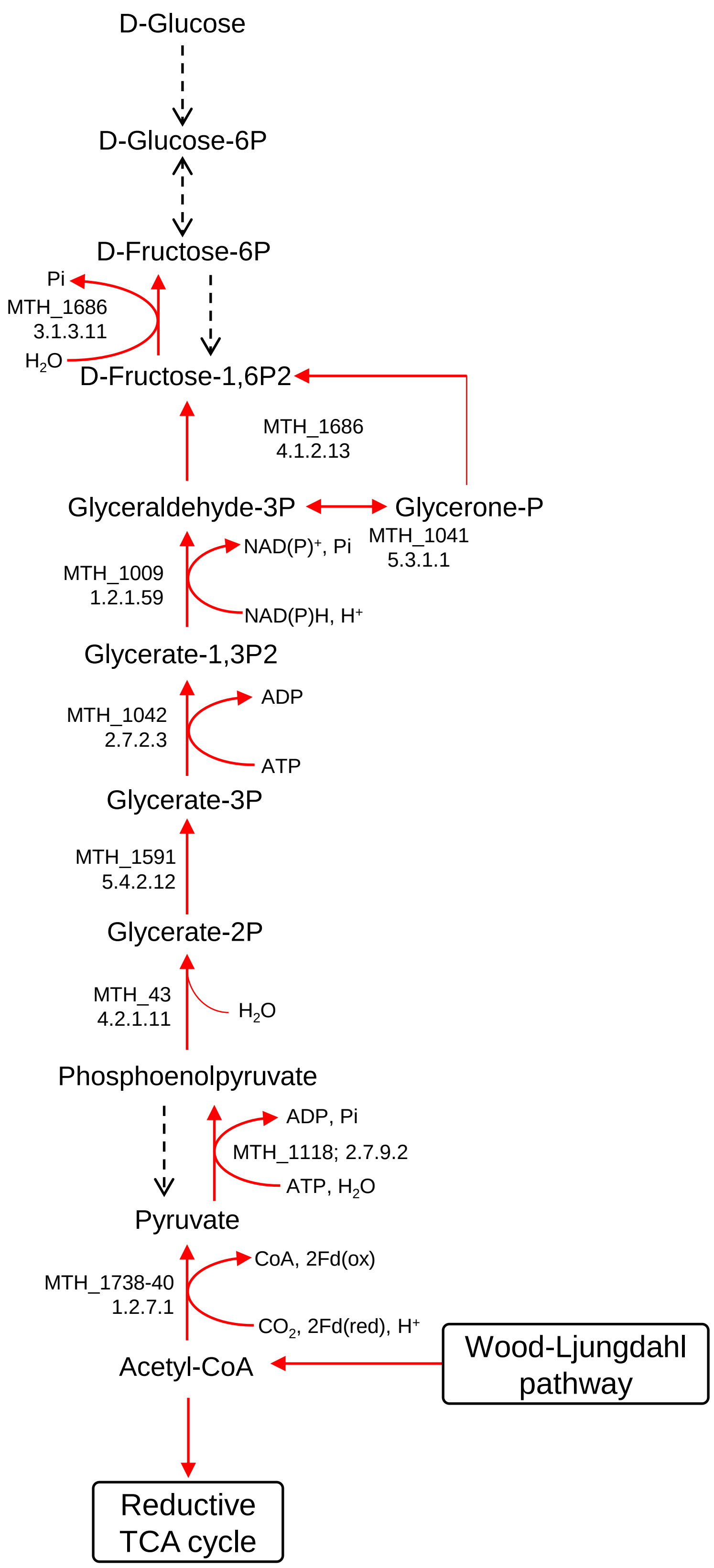

### Wood-Ljungdahl pathway

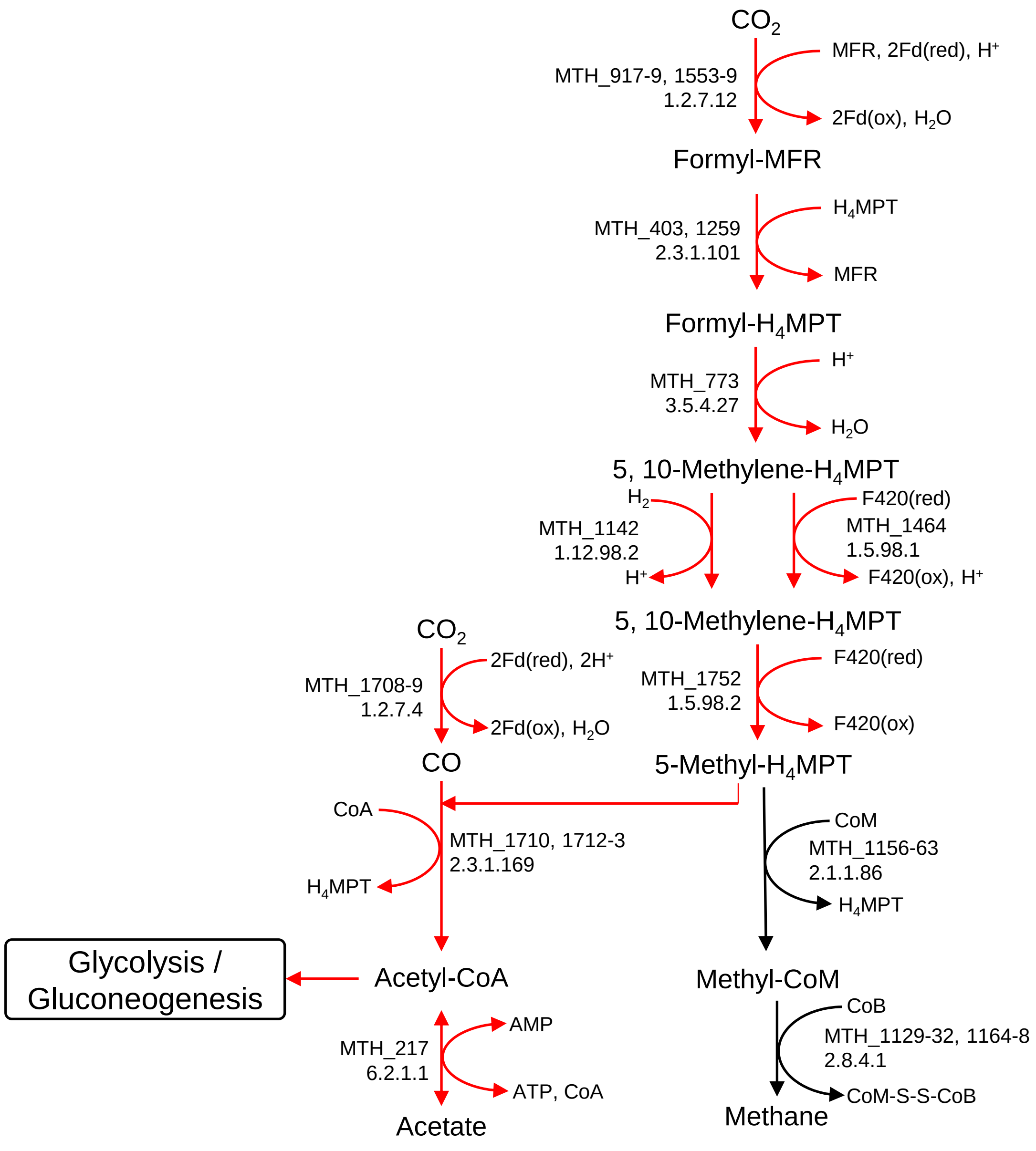

### Reductive TCA cycle

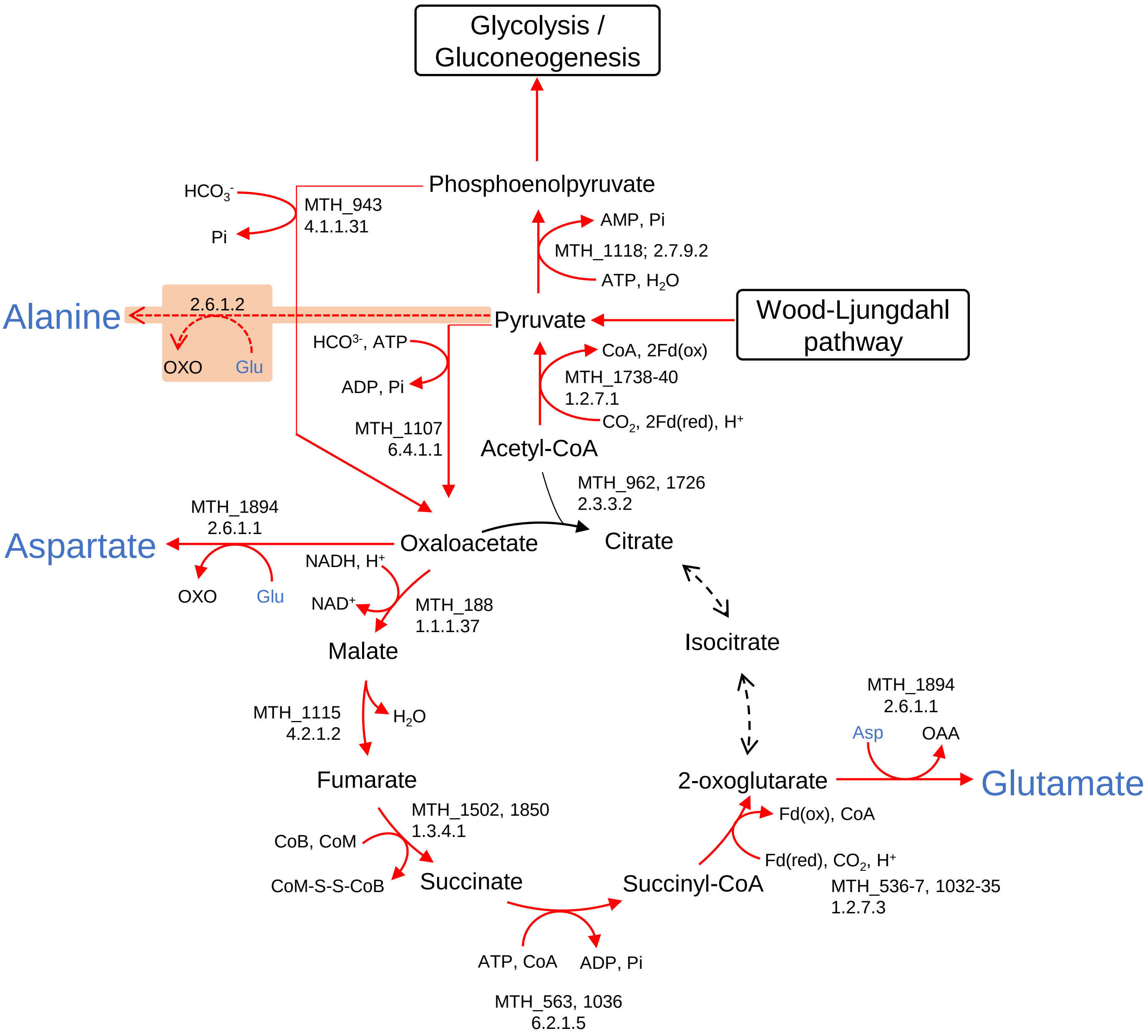

### Glutamine, arginine, and proline biosynthesis pathway

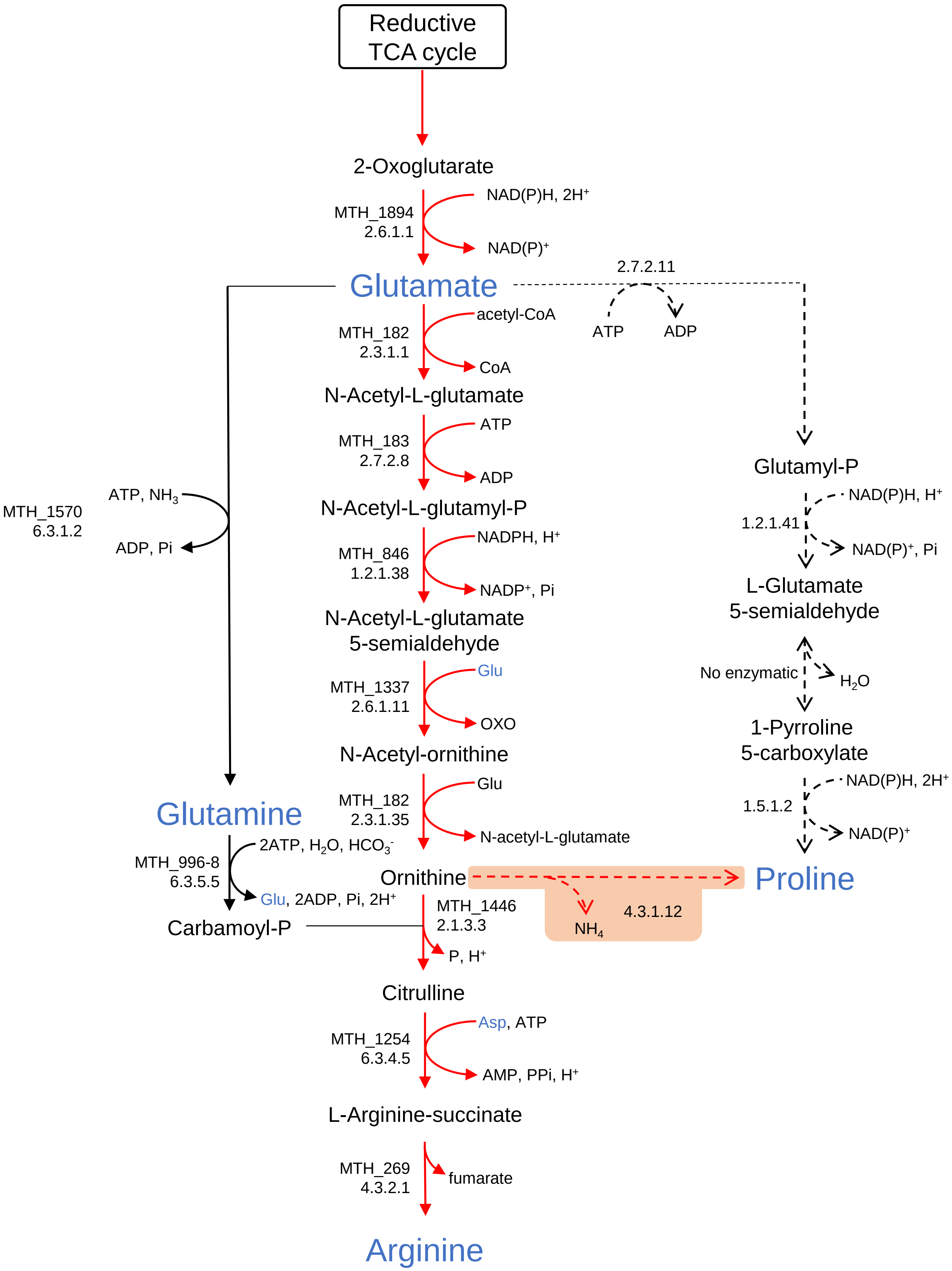

### Glycine, serine, and cysteine biosynthesis pathway

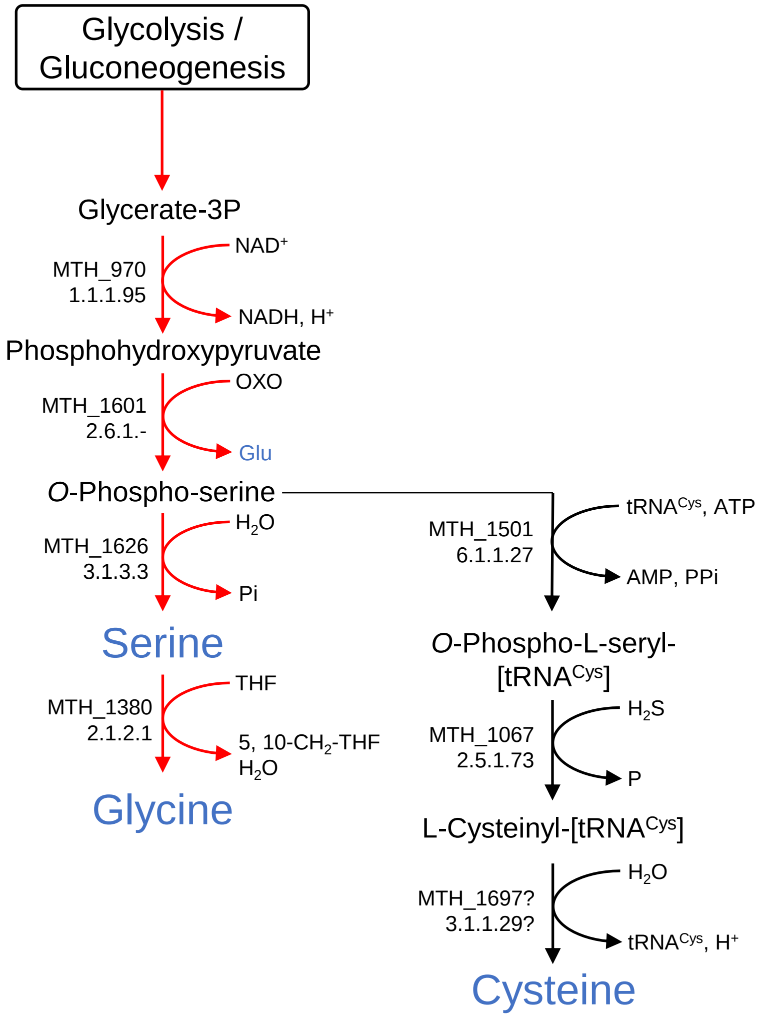

### Lysine, threonine, methionine, and asparagine biosynthesis pathway

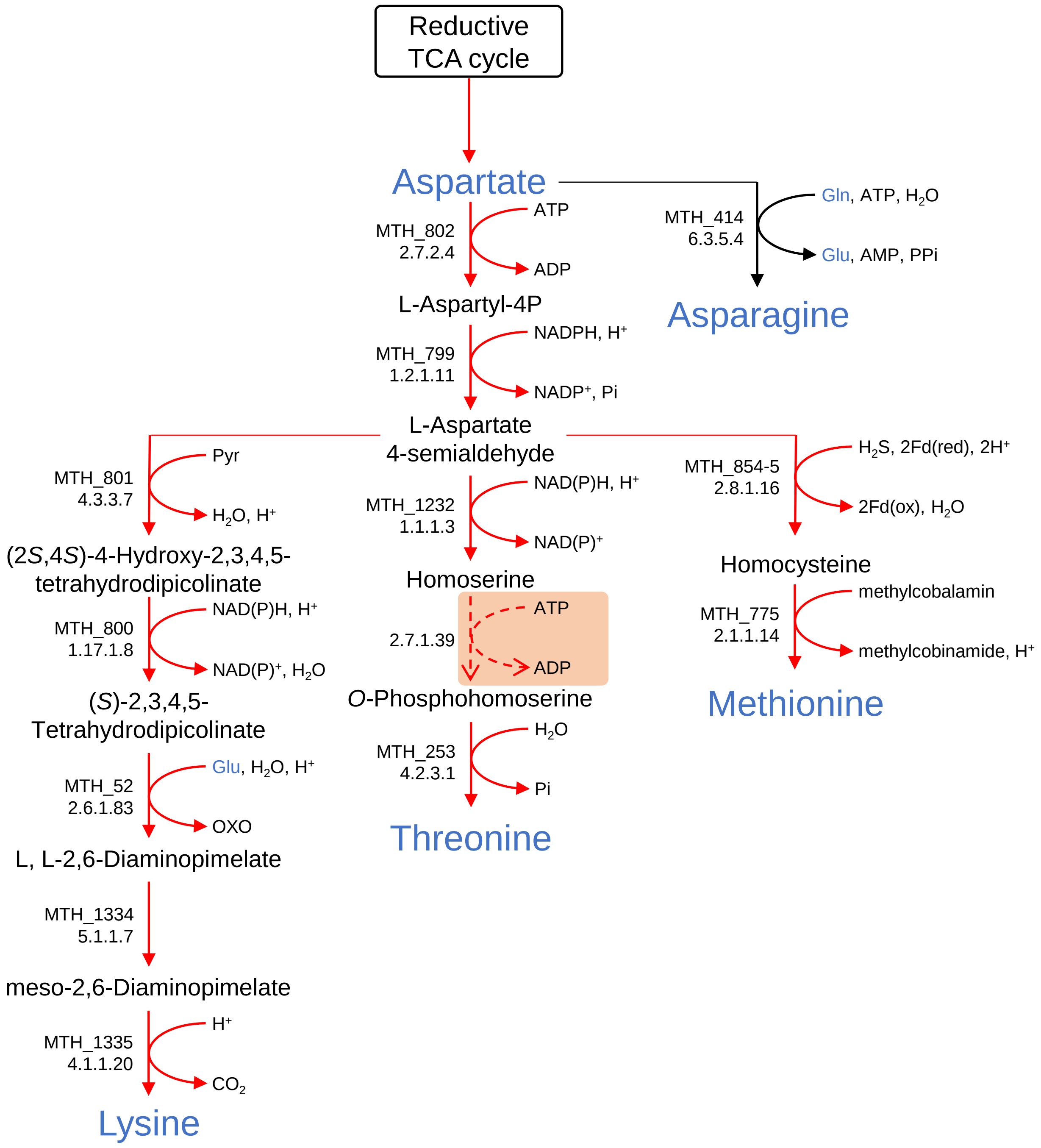

### Histidine biosynthesis pathway

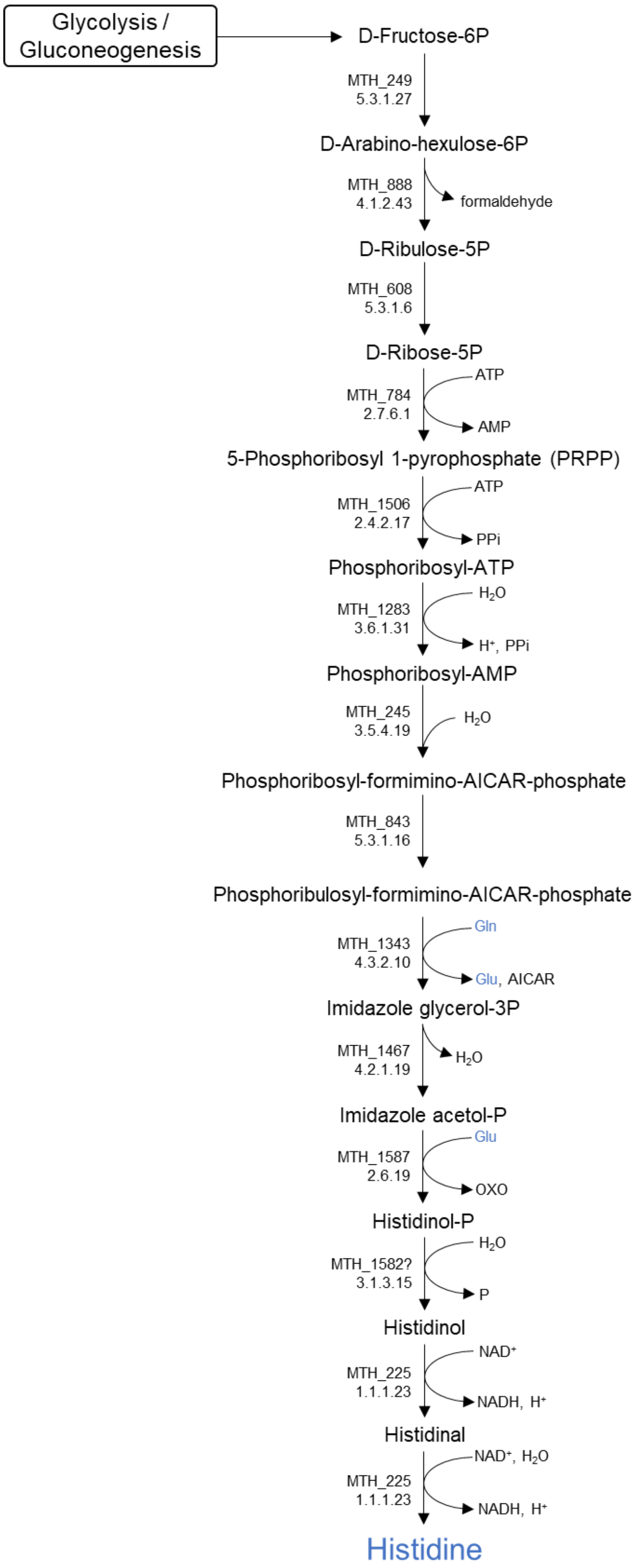

### Tryptophan, phenylalanine, and tyrosine biosynthesis pathway

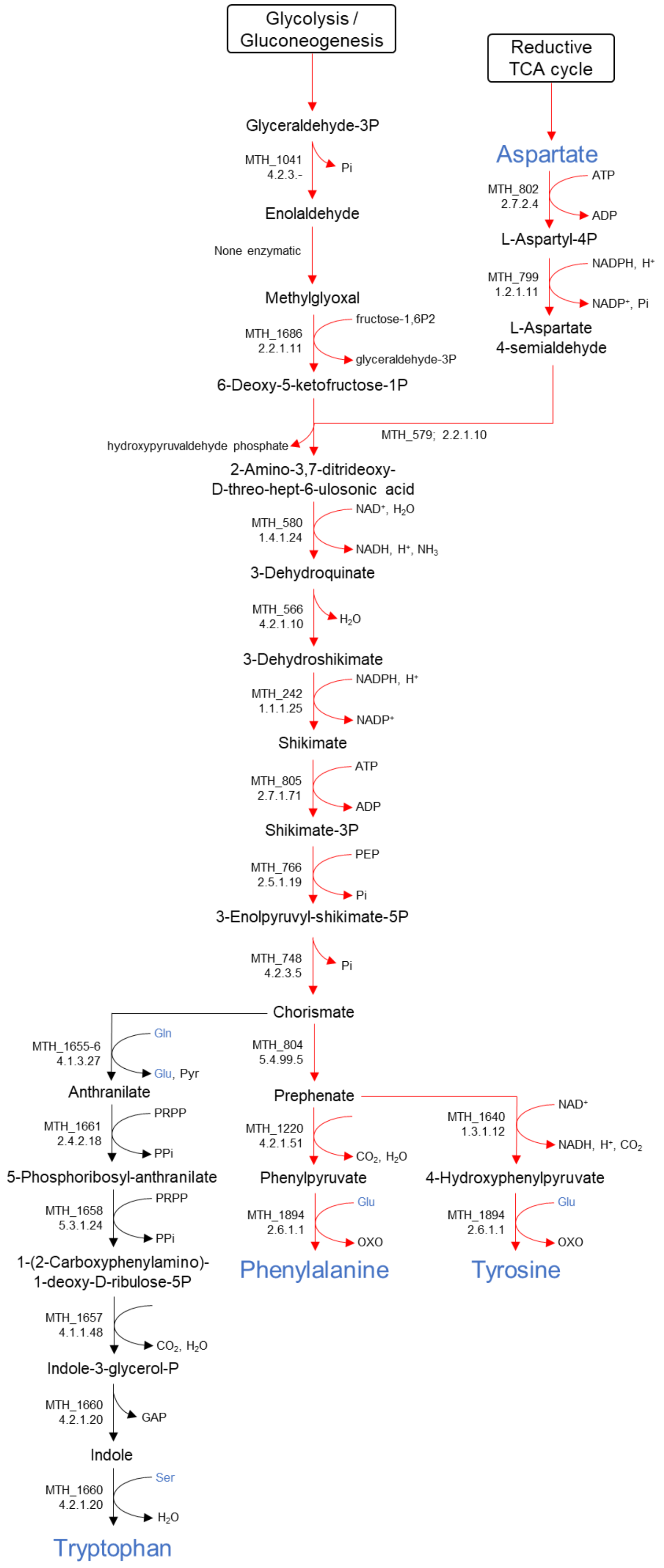

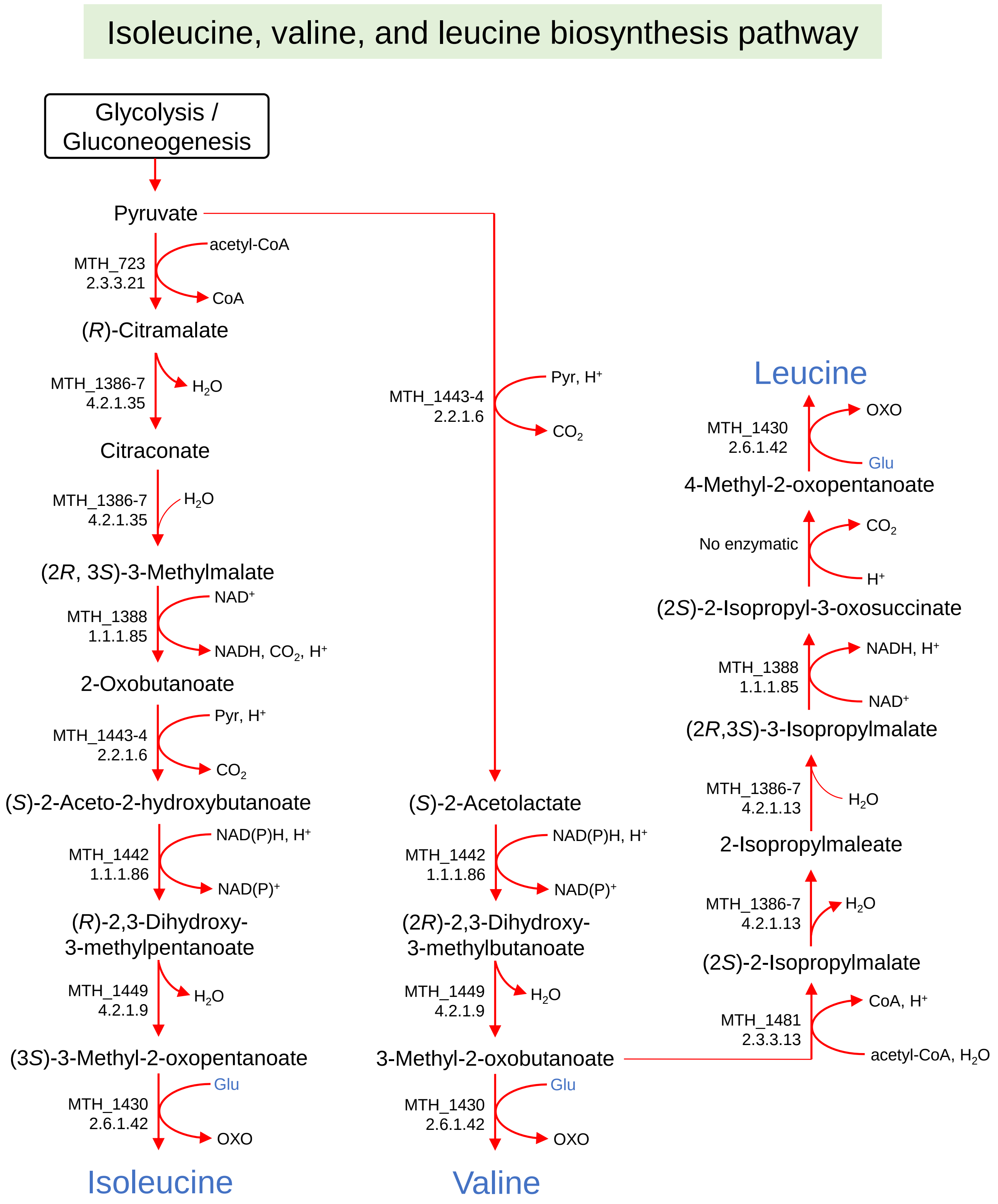


**Fig. S1 Central carbon metabolism and amino acid biosynthetic pathway of *M. thermautotrophicus* under autotrophic growth condition reconstructed from the genome information and isotopomer analysis using the developed CE-MS and CE-MS/MS techniques**

Compounds colored in blue indicate proteinogenic amino acids. Red lines indicate active pathways under autotrophic growth condition confirmed by isotopomer analysis. Dot lines indicate there are no corresponding enzymes found in the genome. The pathways predicted to be present using the tracer-based metabolomics are highlighted in orange. Among 20 proteinogenic amino acids, biosynthetic pathways of asparagine, glutamine, tryptophan, cysteine, and histidine are not conformed in this study.
